# The kinetic Ising model encapsulates essential dynamics of land pattern change

**DOI:** 10.1101/2023.07.04.547706

**Authors:** Tomasz F. Stepinski, Jakub Nowosad

## Abstract

A land pattern change represents a globally significant trend with implications for the environment, climate, and societal well-being. While various methods have been developed to predict land change, our understanding of the underlying change processes remains inadequate. To address this issue, we investigate the suitability of the 2D kinetic Ising model (IM), an idealized model from statistical mechanics, for simulating land change dynamics. We test the IM on a variety of diverse thematic contexts. Specifically, we investigate four sites characterized by distinct patterns, presumably driven by different physical processes. Each site is observed on eight occasions between 2001 and 2019. Given the observed pattern at times *t*_*i*_, *i* = 1, …, 7, we find two parameters of the IM such that the model-evolved land pattern at *t*_*i*+1_ resembles the observed land pattern at that time. Our findings indicate that the IM produces approximate matches to the observed patterns in terms of layout, composition, texture, and patch size distributions. Notably, the IM simulations even achieve a high degree of cell-scale pattern accuracy in two of the sites. Nevertheless, the IM has certain limitations, including its inability to model linear features, account for the formation of new large patches, and handle pattern shifts.

## 1. Intro

Land change studies have gained significant attention due to the accelerated transformation of Earth’s land compared to previous years (Winkler et al., 2021). Over the past few decades, remote sensing of the land surface from space has provided insights into this global environmental trend (Song et al., 2018), which affects a vast majority of landmasses and various land themes. Among these themes, tropical forest deforestation stands out as the most profoundly impacted (Hansen et al., 2013).

The acceleration of land change can be attributed, directly or indirectly, to anthropogenic impacts (Venter et al., 2016). Consequently, global society holds the potential to intervene and mitigate, or even reverse, this trend. However, effective action requires accurate assessments of observed changes and reliable predictions of future changes. To address these requirements, a multitude of methods have been developed to assess the magnitude of past land change (Li et al., 2018; Nowosad et al., 2019) and predict future changes (Veldkamp and Lambin, 2001; Liu and Yang, 2015). The majority of these prediction methods adopt an empirical approach, extrapolating change patterns observed in the past while assuming constraints on future rates of change (Camacho Olmedo et al., 2018). However, this research approach does not establish a causal link between the driving factors (causes) and the resulting patterns of land change (effects).

Causality is of interest in the field of landscape ecology, where the cause and effect are often referred to as the process (the forces acting on the land) and the pattern (the resulting landscape mosaic influenced by these forces) (Turner, 1989). This necessitates the use of deterministic or agent-based modeling approaches. Deterministic models (Rastetter et al., 2003; Scheller et al., 2007; Gustafson, 2013) simulate the impacts of environmental and anthropogenic processes on land patterns through mathematical descriptions of actual processes. On the other hand, agent-based models (Valbuena et al., 2010) simulate the behavior of individual agents (e.g., farmers, developers, or land managers) who interact with each other and their environment, resulting in changes to land patterns. However, a challenge associated with both deterministic and agent-based models lies in accounting for the multitude of potential forces and their interactions. Consequently, constructing and testing such models becomes a complex task due to the large number of free parameters involved.

To facilitate progress, a causal model can be substituted with an idealized model that retains simplicity for analysis or simulation purposes while capturing the fundamental aspects of the observed phenomenon. Within the realm of ecology, these models are known as neutral models of land change (Gardner et al., 1987; Gaucherel et al., 2014; Gaucherel and Houet, 2009). Neutral models are typically stochastic in nature, where the resulting land pattern emerges from the collective influence of random microscopic processes.

This paper aims to assess the applicability of the Ising model (IM) (Ising, 1924; Onsager, 1944; Brush, 1967; Cipra, 1987), a neutral model, as a tool for simulating land change. In this context, the term “land” refers to a pattern of land cover types assigned to cells, which are the smallest square-shaped plots of land arranged in a 2D grid. A “site” represents a specific tract of land corresponding to the entire grid. “Land change” specifically denotes alterations in the pattern of a site over time. It is important to note that our concept of land change aligns with the remote sensing notion of land use/land cover (LULC) change (Cihlar, 2000).

The IM, initially introduced as a model for magnetic substances, has found applications beyond physics, extending into disciplines such as social science (Bornholdt and Wagner, 2002; Stauffer, 2008), psychology (Brandt and Sleegers, 2021; Cramer et al., 2016), genetics (Majewski et al., 2001), environment (Ma et al., 2019), and, notably for this paper, ecology. In the field of ecology, the IM has been employed to investigate various phenomena, including the study of forest canopy-gap structure (Katori et al., 1998; Kizaki and Katori, 1999; Schlicht and Iwasa, 2006), modeling vegetation patterns along a regional rainfall gradient in southern Africa (Scanlon et al., 2007), analyzing population dynamics (Nareddy et al., 2020), and elucidating emergent behaviors like masting (Noble et al., 2018).

In a recent study by Stepinski (2023), the Ising model (IM) was examined as a model for the transition from completely forested to fully agricultural land. However, it is important to note that this model was not explicitly kinetic, focused solely on one thematic context, and assessed the model’s time series using reconstructed patterns from multiple sites rather than using a time series of landscapes from a single site.

Our objective is to explore the suitability of the IM as a simplified representation of various real-life processes responsible for binary land change. Although focusing on binary patterns may appear restrictive, this choice is driven by the capabilities of the IM and the specific interests within the field. Many studies in land use/land cover change (LULC) analysis involve examining the changes within a particular LULC category, such as deforestation (Jamaludin et al., 2022), urbanization (Chen et al., 2020), desertification (Tomasella et al., 2018), or wetland loss (Jamal and Ahmad, 2020). In such applications, the focus is typically on two categories: the foreground, representing the category under investigation for its pattern change, and the background.

It is essential to note that our investigation does not aim to utilize the IM as a tool for predicting future patterns with high cell-level accuracy, nor do we consider it a competitor to empirical predictors. Instead, our focus is on evaluating the feasibility of the IM as a basic dynamic process for simulating land change. While the simulated patterns need to match observations, this matching does not necessarily require high cell-level accuracy.

The Ising model (IM) itself does not inherently possess any predefined dynamics. Therefore, previous applications of the IM in ecology (as mentioned in the references above) have mainly focused on utilizing the IM to simulate equilibrium (steady-state) land patterns and comparing them with observations obtained at a single point in time. However, real land patterns do not exist in a steady state. Multiple observations over time demonstrate that land patterns undergo changes on various time scales, influenced by spatial scale and thematic content (Nowosad et al., 2019). What sets our approach apart is the utilization of a kinetic IM, which enables the simulation of a time-dependent evolution of land patterns. The kinetic IM refers to the IM with a simply-defined temporal evolution rule incorporated.

In order to evaluate the effectiveness of the IM in simulating the evolution of binary land patterns, we selected four specific sites that have undergone land cover changes associated with the loss or gain of distinct land use/cover categories. These categories include forest (reforestation), crops (expansion of croplands), wetlands (loss of wetland), and barren land (expansion of open-pit mining). The data for these sites were obtained from the National Land Cover Dataset 2019 (NLCD2019) (Dewitz, 2021). The NLCD2019 provides maps of sixteen land cover categories for the conterminous United States at a 30-meter resolution for the years 2001, 2004, 2006, 2008, 2011, 2013, 2016, and 2019. By utilizing our IM-based simulator, we were able to identify the best-fit time series of simulated patterns for these selected sites and subsequently compared them to the corresponding observed patterns.

## 2. Model description

The IM is grounded in the principles of statistical mechanics and energy minimization. In the IM, a site is represented by a rectangular array of cells with dimensions *k*_1_ × *k*_2_. For the sake of simplicity, we assume square sites in this paper, hence *k*_1_ = *k*_2_ = *k* and the total number of cells is denoted as *n* = *k*^2^. Each cell in the IM is assigned to one of two categories: cells corresponding to the focus category of land use/land cover (LULC) are assigned a value of 1, while cells representing the background category are assigned a value of - 1. In our figures, we consistently depict focus cells as green and background cells as yellow.

The IM assumes that a cell interacts solely with its four nearest neighbors, following the von Neumann neighborhood scheme. Additionally, the cell is influenced by an external force. The configuration of the array, denoted as ω_*s*_, represents the land’s pattern. Each specific pattern is associated with an energy value, denoted as *E*(ω_*s*_),

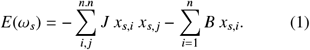

In Eq. (1), *x*_*s,i*_ represents the category of the *i*th cell in pattern *s*. The first term on the right side of Eq. (1) corresponds to an interaction term, with the symbol *n, n* indicating summation over all nearest neighbors in the array. This term captures the propensity for neighboring cells to belong to the same category. The dimensionless parameter *J* denotes the strength of this propensity. In the context of land science, the first term accounts for a degree of the spatial autocorrelation of the landscape.

The second term on the right side of Eq. (1) represents the response of a cell to an externally imposed force, favoring either the focus category (with a positive value of *B*) or the background category (with a negative value of *B*). The dimensionless parameter *B* quantifies the strength of this external force. It is important to note that *x, J*, and *B* are all dimensionless. Consequently, *E*(ω_*s*_) is a dimensionless fitness function, referred to as “energy” for historical reasons. The parameters *B* and *J* are assumed to have uniform values across the entire array.

The original IM includes a third parameter known as temperature, denoted as *T*. However, in numerous non-physics applications of the IM, specifying an equivalent temperature parameter is often challenging. In these cases, the temperature can be understood as representing the willingness or ability of the pattern to deviate from its lowest energy state, potentially accounting for environmental noise (Stauffer and Solomon, 2007). In our study, we assume the temperature to be an unspecified constant that is incorporated into the definitions of *J, B*, and *E*.

Spatial autocorrelation-causing force and external force are generic concepts that can be interpreted differently depending on the thematic context. The spatial autocorrelation process can manifest in various ways (Koenig, 1999). For instance, (a) land-use decisions, such as converting natural vegetation to croplands or urban areas, often lead to autocorrelation due to logistical considerations. (b) Spatial diffusion, such as the spread of invasive species or diseases from one area to another, can also contribute to autocorrelation. (c) Spatial feedback mechanisms, such as the interaction between neighboring ecosystems, can further influence autocorrelation patterns (Fajardo and Velázquez, 2021). Similarly, a category-favoring force can arise through different mechanisms. Examples include (a) the influence of physical and environmental factors on land cover, (b) the impact of business or political decisions on land-use patterns, and (c) land conservation or restoration policies shaping the distribution of land cover categories. The specific nature of the force depends on the context and underlying factors driving land change.

For a given set of values for *J* and *B*, the initial pattern undergoes evolution towards the configuration that minimizes *E*(ω_*s*_). However, as mentioned in the introduction, the IM itself does not provide a description of how this evolution occurs, necessitating the inclusion of a temporal evolution rule. In the literature, three different temporal evolution rules have been proposed for the IM: the Metropolis dynamics (Metropolis et al., 1953), the Glauber dynamics (Glauber, 1963), and probabilistic cellular automata (PCA) dynamics (Procacci et al., 2016; D’Autilia et al., 2021). In this study, we employ the single-flip Glauber dynamics with periodic boundary conditions to simulate land change.

Under the single-flip dynamics, each dynamic step involves an attempt to change the value of a randomly selected cell to its opposite value, essentially flipping it. The success or failure of this attempt depends on the flip probability, which is given by:

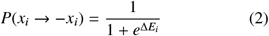

where Δ*E*_*i*_ is,

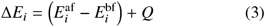

The superscripts “af” and “bf” indicate “before attempted flip” and “after attempted flip,” respectively.

The probability of a flip is 0.5 when Δ*E*_*i*_ = 0, it is high when Δ*E*_*i*_ < 0, and it is low when Δ*E*_*i*_ > 0.

In Eq. (3), the first term on the right-hand side represents the difference between the values of *E*_*i*_ associated with cell *i* after and before the attempted flip. The second term, denoted as *Q*, is not part of the Glauber dynamics or the original IM. It represents our modification of the algorithm aimed at reducing the occurrence of salt-and-pepper noise in the focus category when simulating the evolution of a coarse-textured pattern.

The value of *Q* is zero except in cases where cell *i* is equal to -1 (representing the background category) and all its neighboring cells are also equal to -1. In such cases, with *Q* = 0, the probability of the cell flipping and becoming a focus cell is small but not small enough to prevent the generation of very small patches of the focus category, which is not observed in reality. By setting *Q* ≫ 0, this probability becomes negligible, effectively eliminating the generation of small patches and eliminating the salt-and-pepper noise in the focus category.

For a cell with *Q* = 0, the change in energy Δ*E*_*i*_ is determined by the values of the cell *i* before and after an attempted flip, the sum *S* _*i*_ of neighboring cell values (cells in the von Neumann neighborhood of the central cell), and the parameters *J* and *B*:

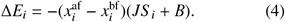

The quantity *S* _*i*_ can take five possible values: -4, -2, 0, 2, and 4. On the other hand, the quantity 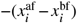 has three possible values: 2, 0, and -2, where the value of 0 corresponds to no flip.

Fig. 1 illustrates ten possible types of von Neumann neighborhoods. Each neighborhood’s center cell has a high probability of flipping if the condition below the neighborhood is true. For the center neighborhood in each row and in the absence of pressure to autocorrelate (*S* _*i*_ = 0), a high flip probability requires a small external push (*B*) towards the opposite category of the central cell. For the two leftmost neighborhoods in the top row, the flip probabilities can be high even in the presence of an external push in favor of the green category (positive *B*), as long as *B* has a sufficiently low absolute value. On the other hand, in the two rightmost neighborhoods in the top row, the flip probability can be high only if there is an external push in favor of the yellow category (negative *B*) with a high enough absolute value. The discussion of the bottom row in Figure 1 is analogous.

**Figure 1:**
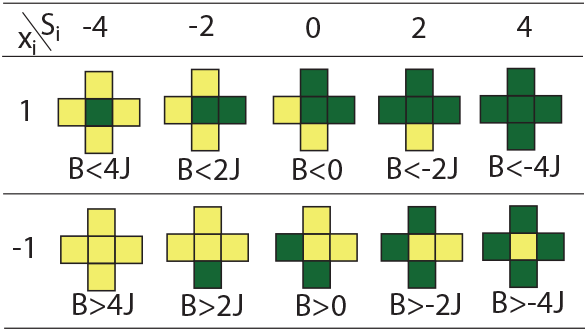
Ten possible types of von Neumann neighborhoods. If a condition below a neighborhood is met, the central cell has a high probability of flipping.

### 2.1 Time unit

We aim to simulate a land change in a way depicted in Fig. 2A. The site shown in Fig. 2A undergoes deforestation, and it has been observed and mapped at eight different time instances. The time intervals between consecutive observations, denoted as (Δ*t*)_*i*_, where *i* = 1, …, 7, are not constant. At each time interval, the area covered by the focus category (green) decreases. However, it is important to note that the rate of this decrease is not constant, as illustrated in Fig. 2B.

**Figure 2:**
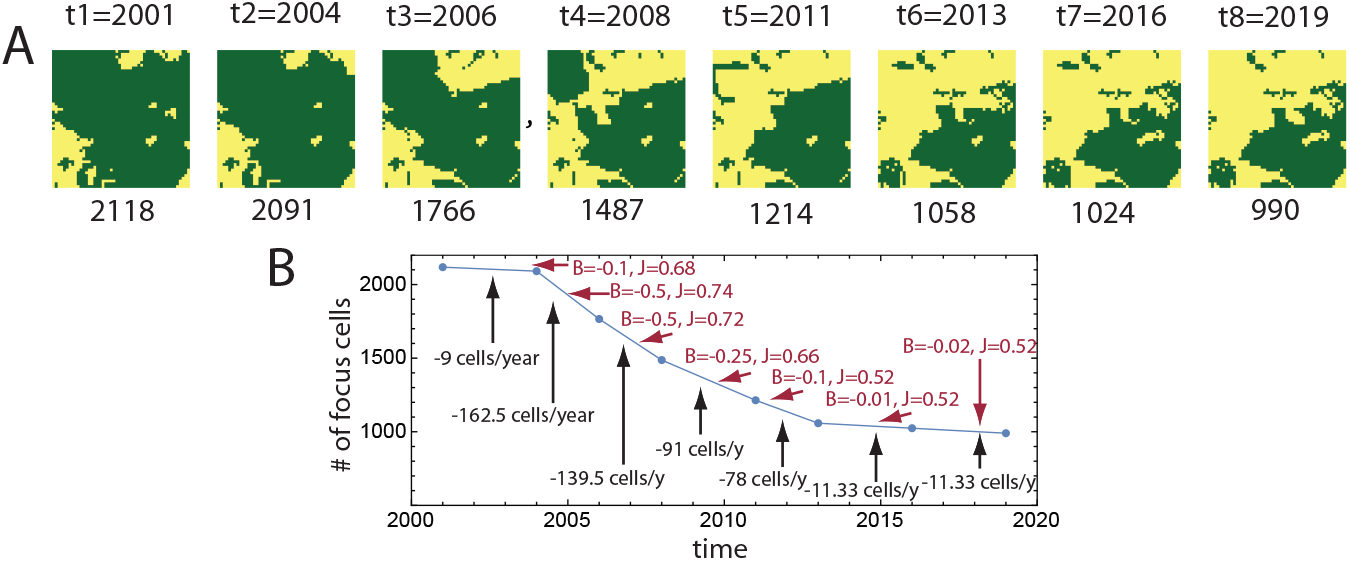
(A) The observed evolution of the land cover pattern from 2001 to 2019 is depicted for a small site measuring 1.5 km × 1.5 km, equivalent to 2,500 cells. The focus category in this case is forest, and the observed process is deforestation. The numbers below the patterns indicate the count of focus cells. (B) A graph is presented to illustrate the temporal loss of focus (forest) cover during the seven transitions. The black descriptors represent the observed loss rates in cells per year, while the red descriptors represent the calculated values of the best-fit parameters for the simulated process.

Our objective is to simulate the observed time series of patterns using the IM, with a focus on determining the optimal values for the IM parameters (*B* and *J*) for each transition in the series. In other words, we aim to find the best-fit values of *B* and *J* that result in the most accurate simulation of pattern changes during each time period (Δ*t*)_*i*_. In Fig. 2B, these best-fit parameter values are indicated in red. To obtain these values, we initiate the IM simulation with the observed pattern at time *t*_*i*_ (serving as the initial condition) and perform a series of Glauber dynamic steps corresponding to (Δ*t*)_*i*_. The goal is to generate a simulated pattern at *t*_*i*+1_ that closely resembles the observed pattern at that time.

To ensure that the number of dynamic steps taken is proportional to (Δ*t*)_*i*_, it is necessary to establish a unit of time that corresponds to the duration of a dynamic step. For instance, in the case of simulating the transition from 2001 to 2004 (as depicted in Fig. 2A), if we use 30,000 dynamic steps, we can set the time unit of a single dynamic step to 3 years divided by 30,000, which equals 0.876 hours. This time unit can be interpreted as follows: in the landscape represented in Fig. 2A, a random cell within the array has an opportunity to change its land cover category every 0.876 hours. The magnitude of this opportunity depends on the parameters of the IM model and the current configuration of the landscape pattern.

Importantly, using a specific time for the dynamic step allows us to determine the number of steps required during the simulation of each transition. For the consecutive transitions shown in Fig. 1A the number of steps is 30,000, 20,000, 20,000, 30,000, 20,000, 30,000, and 30,000. A different time unit can be either larger or smaller than 0.876 hours, but it must remain consistent throughout the simulation to ensure that the number of steps for each transition is proportional to its duration.

### 2.2 Simulation

The simulation is performed on a transition-by-transition basis. Let’s consider a specific transition from time *t*_1_ to time *t*_2_ with a duration of (Δ*t*)_1_ years. In our simulation, we employ *n* dynamic steps per year, where *n* represents the number of cells in the array. Theoretically, this means that each cell has the opportunity to undergo a flip once per year.

The similarity between the observed and simulated patterns at time *t*_*i*+1_ is quantified using the Euclidean distance between the observed and simulated data points, denoted as (*m*, ⟨*s*_*i*_ *s* _*j*_⟩)_obs_ and (*m*, ⟨*s*_*i*_ *s* _*j*_⟩)_sim_, respectively. Here, *m* represents the composition imbalance index of the landscape, which is calculated as *m* = 2 *f* − 1, where *f* is the fraction of focus cells in the site. The range of *m* lies between -1 (indicating only background cells) and 1 (indicating only focus cells).

Additionally, ⟨*s*_*i*_ *s* _*j*_⟩ is a measure of the landscape’s spatial autocorrelation, referred to as the texture index. It ranges from 0 (indicating fine texture) to 1 (indicating coarse texture). Our objective is to determine the values of *B* and *J* that yield the highest similarity between the observed pattern at *t*_*i*_ and the simulated pattern at *t*_*i*+1_, while considering the constraints of the number of dynamic steps (proportional to (Δ*t*)_*i*_) and periodic boundary conditions.

The best-fit values of *B* and *J* are determined using the simulated annealing optimization algorithm (Kirkpatrick et al., 1983). We employ the implementation of this algorithm provided by the optimization package in R (Husmann et al., 2017). A detailed example of the optimization workflow can be found in the vignette of the spatialising R package (Nowosad, 2023).

Since the Glauber dynamics is stochastic in nature, we repeat the aforementioned procedure 200 times to obtain an ensemble of best-fit parameter pairs (*B* and *J*). From this ensemble, we select the pair that corresponds to the peak of the frequency distribution of pairs. This chosen pair represents our final solution for the parameters, characterizing the magnitude and nature of the process governing the pattern change during the transition from *t*_*i*_ to *t*_*i*+1_. The same procedure is applied to determine the solutions for the remaining transitions.

## 3. Results

To assess the feasibility of the IM to simulate the land change we conducted calculations for four sites. The sites were selected to exhibit different processes leading to the change of pattern with time, reforestation, expansion of croplands, wetland loss, and open-pit mining. The observed change is documented by the NLCD2019 dataset that shows LULC maps of those sites at eight different times starting in 2001 and ending in 2019. Each site is represented by a time series of eight arrays of *n* = 250 × 250 = 62,500 LULC-labeled cells. Calculations are conducted using a protocol as described in section 2.

Best-fit values of parameters *B* and *J* for each transition are shown in Table 1. The simulated land’s pattern is compared to an observed land’s pattern in multiple ways. First, we compare values of *m* (expressed in terms of the number of focus cells) and ⟨*s*_*i*_ *s* _*j*_⟩ in simulated and observed land patterns. The similarity of these values indicates a similarity in the area and texture of the focus cells’ observed and simulated patterns.

**Table 1:** Best-fit process parameters

| process | coordinates | 2001→04 | 2004→06 | 2006→08 | 2008→11 | 2011→13 | 2013→16 | 2016→19 |
| --- | --- | --- | --- | --- | --- | --- | --- | --- |
| reforestation<br>Maine | 69.31298 W<br>46.65407 N | B=0.18<br>J=0.42 | B=0.48<br>J=0.40 | B=0.76<br>J=0.38 | B=0.05<br>J=0.45 | B=0.09<br>J=0.45 | B=0.095<br>J=0.45 | B=0.06<br>J=0.45 |
| croplands gain<br>Iowa | 93.28743 W<br>40.86183 N | B=0.16<br>J=0.50 | B=0.25<br>J=0.50 | B=0.22<br>J=0.50 | B=0.28<br>J=0.45 | B=0.15<br>J=0.50 | B=0.07<br>J=0.50 | B=-0.001<br>J=0.55 |
| wetland loss<br>Florida | 81.85418 W<br>26.86219 N | B=0.04<br>J=0.34 | B=-0.06<br>J=0.32 | B=-0.03<br>J=0.32 | B=-0.12<br>J=0.33 | B=-0.05<br>J=0.33 | B=-0.03<br>J=0.34 | B=-0.045<br>J=0.35 |
| open-pit mining<br>Wyoming | 105.28693 W<br>43.69249 N | B=0.15<br>J=0.50 | B=0.8<br>J=0.50 | B=0.4<br>J=0.50 | B=0.47<br>J=0.50 | B=-0.03<br>J=0.60 | B=0.732<br>J=0.60 | B=0.28<br>J=0.50 |

Second, we compare mapped and simulated complementary cumulative distribution functions (cCDF) of patch size and area. Patches are sets of adjacent cells of focus category, they are extracted using the connected components labeling algorithm (Rosenfeld and Pfaltz, 1966). The cCDF is the probability that the variable takes a value greater than *x*. For example, in the case of the patch size distribution, cCDF(10)=0.1 means that 10% of patches have sizes larger than 10 cells. In the case of the area distribution cCDF(10)=0.9 means that 90% of the area covered by the focus category is in patches having a size larger than 10 cells.

Third, we treat simulation as a prediction and calculate prediction metrics, accuracy, recall (for the focus category), and precision (for the focus category). A recall is the estimated probability that a cell randomly selected from among focus cells in the observed landscape is also a focus cell in the simulated landscape. Precision is the estimated probability that a cell randomly selected from among focus cells in the simulated pattern is also a focus cell in the observed pattern. High values of recall and precision indicate individual cell-level agreement between two patterns. Top empirical models achieve values of recall and precision of ∼ 90% (see, for example, Kumar and Agrawal (2022).

### 3.1 Reforestation

Our first site is located in Aroostook county in northern Maine. This county has a notable historical trend of deforestation; however, a significant shift has occurred in the forest condition since approximately 2005 (Acheson, 2008). The selected site serves as an illustrative example of this turnaround. The results obtained for this site are presented in Fig. 3, which consists of five rows.

**Figure 3:**
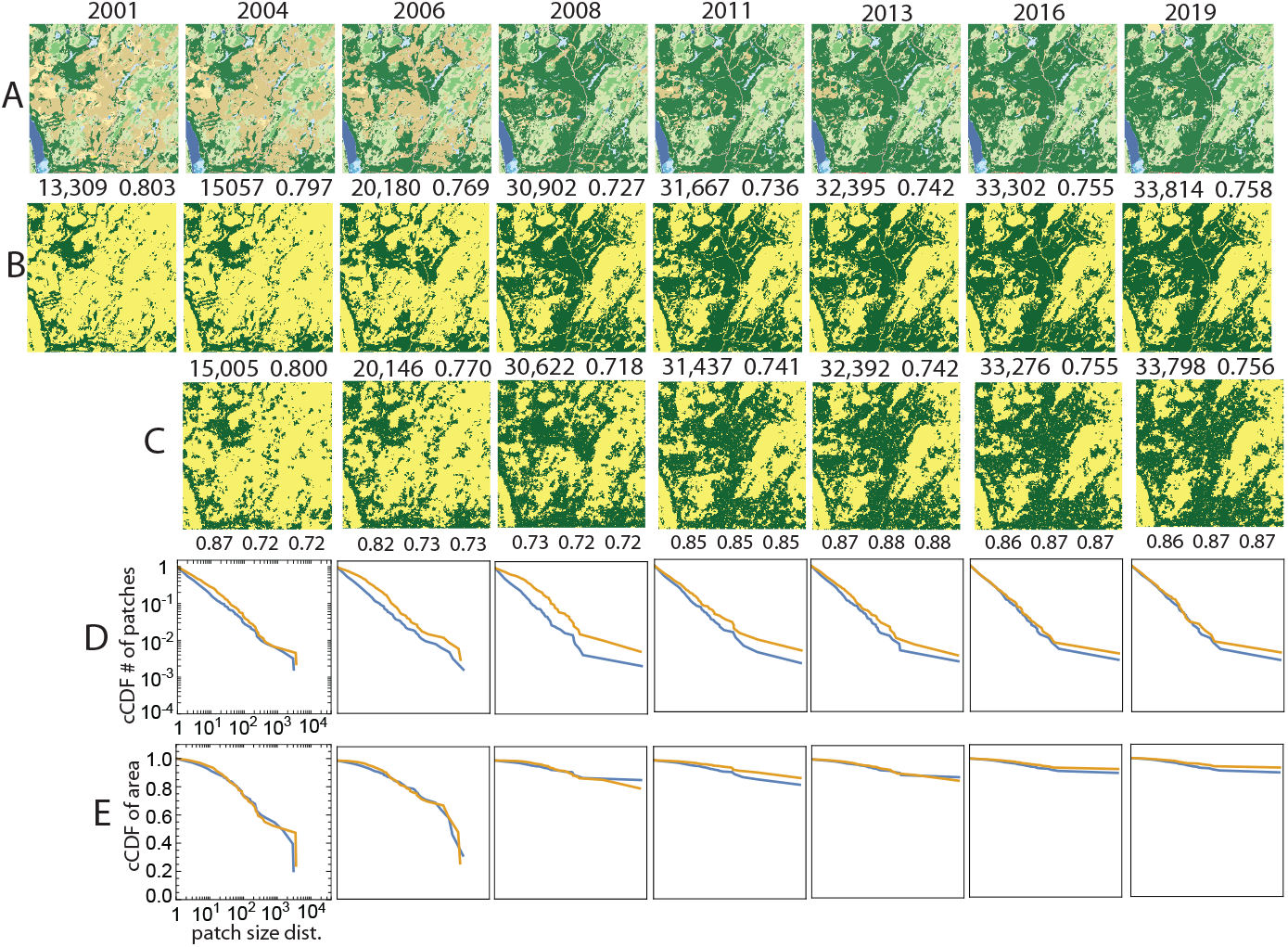
Comparison between the observed and the simulated land change at a site in the US state of Maine undergoing the reforestation process during the 2001-2019 period. (A) The series of original NLCD2019 maps of LULC categories. (B) NLCD2019 maps reclassified into two categories, the focus category (evergreen forest) and the background. (C) Simulated land change. (D) Complementary CDFs of patch size for the patterns in row B (blue) and row C (orange). (E) Complementary CDFs of patch area for the patterns in row B (blue) and row C (orange). The numbers above observed and simulated patterns are the number of forest cells and the pattern’s texture parameter ⟨*s*_*i*_ *s* _*j*_⟩. Numbers below the simulated patterns are accuracy, recall, and precision.

In the first row of Fig. 3, NLCD2019 maps of the site from 2001 to 2019 are displayed, where different colors represent distinct LULC categories. The second row exhibits the same NLCD2019 maps, but this time they are reclassified into binary patterns. In these reclassified patterns, the focus category is represented by dark green color, corresponding to the evergreen forest category (category 42 in the NLCD2019 maps), as depicted in row A of the figure.

The third row of Fig. 3 displays the results of our simulations, utilizing the best-fit values of *B* and *J* for each transition (listed in the first entry of Table 1). Upon visual inspection, the observed time series (row B) and the simulated time series (row C) exhibit a striking similarity. However, upon closer examination, some discrepancies can be observed, particularly during the period of the most significant change, (Δ*t*)_2_ and (Δ*t*)_3_. Quantitatively, both the observed and simulated time series are characterized by nearly identical values of *m* and ⟨*s*_*i*_ *s* _*j*_⟩, indicating a high degree of similarity in terms of com-position and texture. The recall and precision values are approximately 70% from 2004 to 2008, corresponding to a period of rapid change, and approximately 90% from 2011 to 2019, during a period of slower change.

The fourth row of Fiig. 3 illustrates the patch size distributions for both the observed patterns (blue) and the simulated patterns (orange). Overall, the patch size distributions of the observed and simulated patterns exhibit a high degree of similarity. Any differences observed in the size distributions primarily arise from variations in the size and/or number of the largest patches. These differences may not be apparent when examining the patterns themselves since the distinction between a single large patch and multiple smaller patches may depend on the presence or absence of a single cell that connects the larger patches.

The fifth row presents the area distributions for the observed patterns (blue) and the simulated patterns (orange). The area distributions of both the observed and simulated patterns demonstrate remarkable similarity. It is noteworthy that as time progresses, a significant majority of focus cells aggregate into a single, very large patch. This indicates that the initially fragmented forest gradually reconnects, forming a cohesive and connected forest structure.

The values of *B* exhibit temporal variability throughout the period from 2001 to 2019. This variability is strongly correlated with the fluctuation of the reforestation rate, as indicated by a correlation coefficient of 0.99. This finding suggests that an external force is responsible for driving the temporal changes in the reforestation rate. On the other hand, the values of *J* remain relatively constant over the entire 2001-2019 period, with an approximate value of *J* ≈ 0.4. This suggests that the propensity of the land to exhibit spatial auto-correlation remained consistent throughout the studied period. However, the specific mechanisms underlying the external influence and the tendency for autocorrelation are beyond the scope of this paper and will require further investigation.

### 3.2 Expansion of croplands

The expansion of croplands in the United States is causing a decline in grasslands and other ecosystems. One of the regions experiencing significant expansion is southern Iowa, as documented by Lark et al. (2020). Our second site, located at the boundary between Lucas and Wayne counties in Iowa, serves as an illustrative example of this expansion. The results for this site are presented in Fig. 4. The figure follows the same organization as Fig. 3, with the green color in rows B and C indicating cultivated crops (NLCD category 82), represented by a brown color in row A.

**Figure 4:**
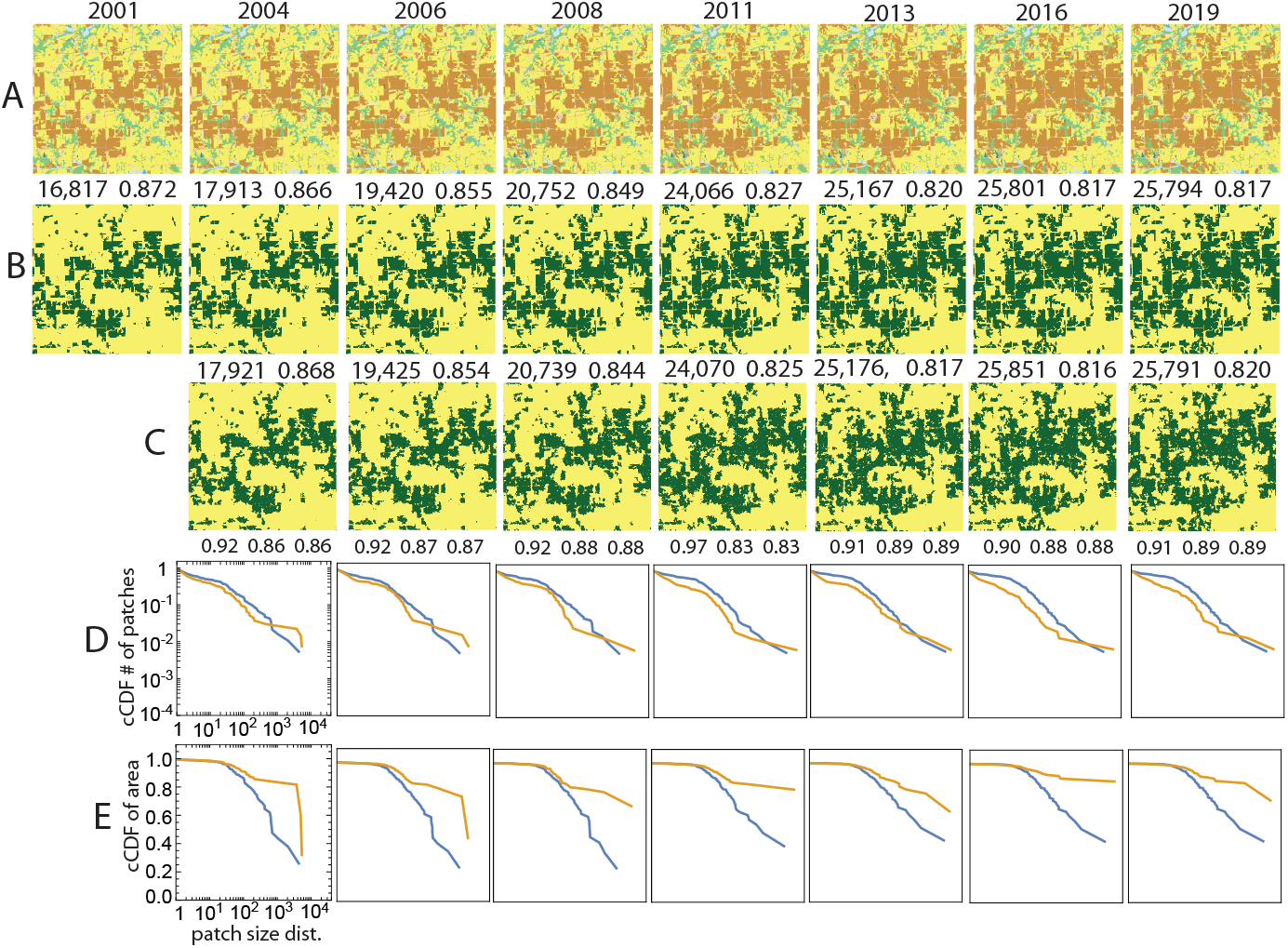
Comparison of the observed and the simulated land change at a site in the US state of Iowa undergoing a crop expansion process during the 2001-2019 period. (A) The series of original NLCD2019 maps of LULC categories. (B) NLCD2019 maps reclassified into two categories, the focus category (crops) and the background. (C) Simulated land change. (D) Complementary CDFs of patch size for the patterns in row B (blue) and row C (orange). (E) Complementary CDFs of patch area for the patterns in row B (blue) and row C (orange). The numbers above observed and simulated patterns are the number of cropland cells and the pattern’s texture parameter ⟨*s*_*i*_ *s* _*j*_⟩. Numbers below the simulated patterns are accuracy, recall, and precision.

The second row in Table 1 presents the best-fit values of *B* and *J* for each transition in the series depicted in Fig. 4. Similar to the reforestation site discussed in subsection 3.1, we observe a high visual similarity between the simulated and observed series. However, there is a notable difference in the presence of grid-like northsouth and east-north features in the observed patterns that are absent in the simulated patterns. These features can be attributed to a historical system dating back to the early days of the United States, where land was divided into one-square-mile quadrangles where feasible. While the IM cannot replicate these grid-like features, it accurately reproduces the overall changing arrangement of croplands in this site.

The quantitative analysis reveals that the simulated and observed patterns exhibit almost identical values of *m* and ⟨*s*_*i*_ *s* _*j*_⟩. It is important to recall that our criterion for determining the best-fit values of *B* and *J* relies on the similarity between the simulated and observed values of *m* and ⟨*s*_*i*_ *s* _*j*_⟩, which serves as the fitness function. The excellent fit obtained indicates that the IM can be effectively tailored to the data. Additionally, the recall and precision values are approximately 70% from 2004 to 2008 (during periods of rapid change) and around 90% from 2011 to 2019 (during periods of slower change). These recall and precision values are comparable to those achieved by empirical models of land change, indicating the effectiveness of the IM in capturing the dynamics of the studied site.

Row D in Fig. 4 illustrates the patch size distributions for the observed patterns (blue) and simulated patterns (orange). It is important to note that the y-axis represents the probability that a randomly selected patch has a size equal to or larger than the corresponding value on the x-axis. Upon examining the series of distributions in row D, we can observe that the observed land patterns exhibit a relatively higher frequency of smaller patches and a relatively lower frequency of larger patches compared to the simulated patterns. This finding is further supported by row E in Fig. 4. Despite the high values of recall and precision, the discrepancies observed in the size and area distributions can be attributed to the IM’s inability to reproduce the linear background features mentioned earlier, which exist in the observed land and contribute to the division of the area into smaller patches.

Similar to the deforestation example discussed in section 3.1, the cropland expansion case also exhibits a strong correlation (0.87) between the inferred values of *B* and the rate of cropland gain. On the other hand, the inferred values of *J* remain relatively constant throughout the 2001-2019 period. This observation leads to a hypothesis that external influences, potentially of an economic nature (Lark et al., 2020), are responsible for the temporal changes in cropland expansion, while the autocorrelation tendency is an inherent characteristic of the site that remains unchanged over the observed time period.

### 3.3 Loss of herbaceous wetlands

Our third site is situated within the Fred C. Babcock/Cecil M. Webb Wildlife Management Area (WMA) in southwestern Florida. This particular site is predominantly composed of two types of wetlands: woody wetlands (NLCD category 90, depicted in a lighter blue shade in Fig. 5A) and herbaceous wetlands (NLCD category 95, depicted in a darker blue shade in Fig. 5A). Our focus category for analysis is the herbaceous wetlands, which exhibits distinct patterns compared to the previous examples, characterized by a finergrained structure.

**Figure 5:**
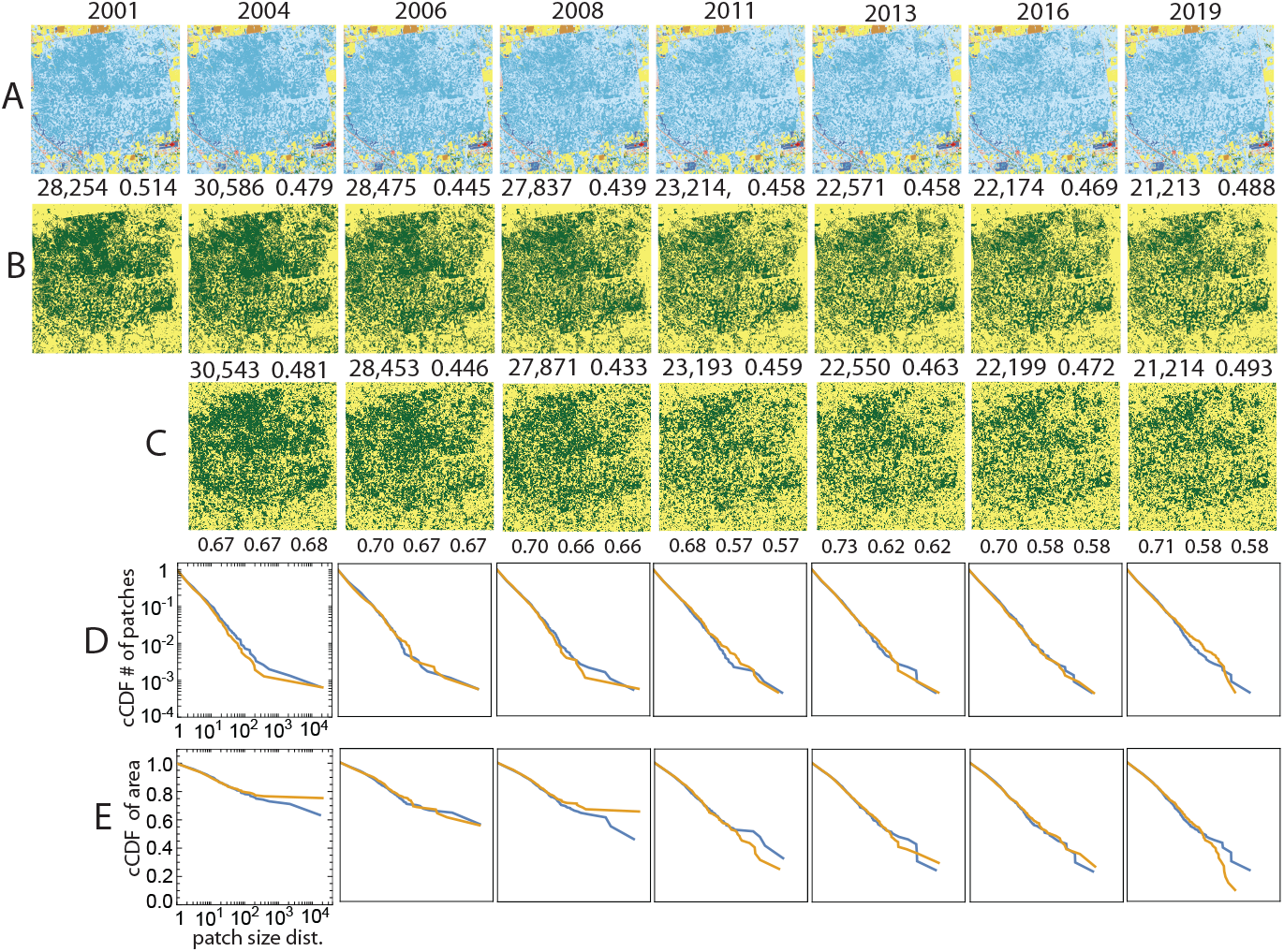
Comparison of the observed and the simulated land change at a site in the US state of Florida undergoing a herbaceous wetlands loss process during the 2001-2019 period. (A) The series of original NLCD2019 maps of LULC categories. (B) NLCD2019 maps reclassified into two categories, the focus category (herbaceous wetlands) and the background. (C) Simulated evolution of land change. (D) Complementary CDFs of patch size for the patterns in row B (blue) and row C (orange). (E) Complementary CDFs of patch area for the patterns in row B (blue) and row C (orange). The numbers above observed and simulated patterns are the number of herbaceous wetland cells and the pattern’s texture parameter ⟨*s*_*i*_ *s* _*j*_⟩. Numbers below the simulated patterns are accuracy, recall, and precision.

The third row in Table 1 presents the best-fit values of *B* and *J* for each transition in the series depicted in Fig. 5. Notably, the values of *B* are negative, indicating a decrease in the area of the focus category. The values of *J* remain relatively constant for the duration of the observed period, but they are smaller compared to the previous two sites, suggesting a lower inherent tendency for spatial autocorrelation. This observation aligns with the fine-grained nature of the landscape pattern observed in this particular site.

Similar to the previous two examples, we observe a strong visual resemblance between the simulated and observed land series in this case. However, it is important to note that visual assessment may not be entirely reliable due to the limited spatial variability in the wetlands patterns at this small scale. To obtain a more accurate evaluation, we rely on quantitative measures. Quantitatively, the simulated and observed patterns exhibit nearly identical values of *m* and ⟨*s*_*i*_ *s* _*j*_⟩, indicating that the Ising model can effectively capture and reproduce this type of pattern as well.

The values of recall and precision in this case are in the range of 70-60%, which is lower compared to the previous two examples. We attribute this lower celllevel accuracy to the fine-grained nature of the wetlands pattern. The stochasticity of the Glauber dynamics in the Ising model leads to distinct realizations of the simulation at the cell level, resulting in differences between individual patterns. This variability is more pronounced in fine-grained patterns, leading to a decrease in accuracy. However, it is worth noting that despite the lower cell-level accuracy, the distributions of patch sizes and areas in both the observed and simulated patterns exhibit a high degree of similarity, as the majority of patches in this landscape are small.

The loss of herbaceous wetlands exhibits a high correlation (0.99) with the inferred values of parameter *B*. This finding aligns with our previous examples and reinforces our conclusion that parameter *B* governs the temporal variability of landscape composition. In this specific case, it influences the loss of herbaceous wetlands. On the other hand, parameter *J* remains consistent and does not undergo significant changes over the observed time scale, highlighting its role as a property inherent to the site.

### 3.4 Open-pit mining

The fourth site encompasses the Black Thunder Coal Mine located in Wyoming. This site is characterized by an open-pit mine, where the landscape predominantly consists of barren land (NLCD category 31), depicted by a gray color in the NLCD maps shown in Fig. 6. Additionally, the site includes grassland (NLCD category 71) depicted by a vanilla color, and shrubland (NLCD category 52) depicted by a beige color. The focus category in this case is the barren land, which corresponds to the pit within the mine. Notably, the pattern evolution in this site deviates from the previous examples, as the pit has undergone a westward shift during the 20012019 period.

**Figure 6:**
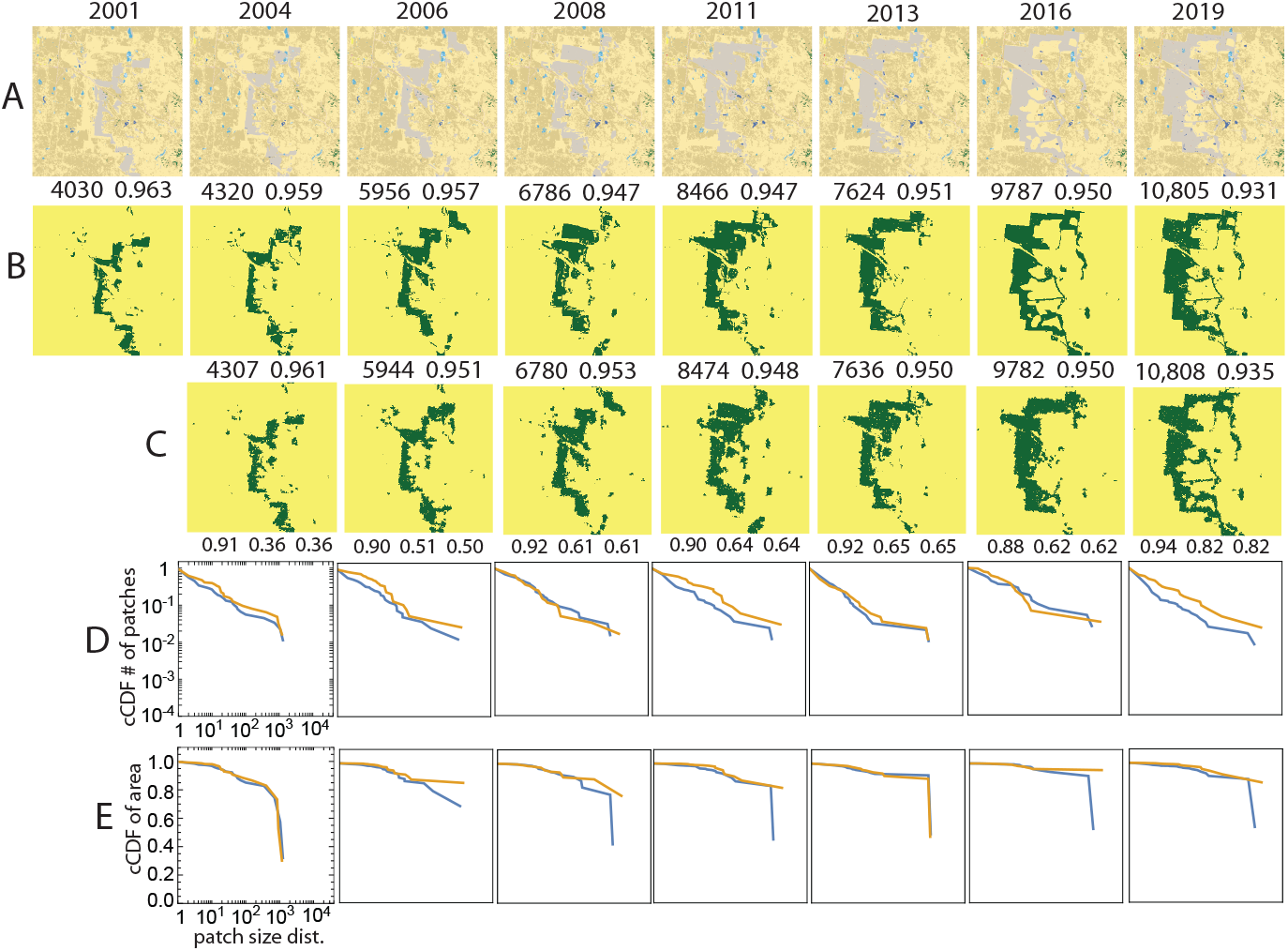
Comparison of the observed and the simulated land change at a site in the US state of Wyoming undergoing a change due to mining activity during the 2001-2019 period. (A) The series of original NLCD2019 maps of LULC categories. (B) NLCD2019 maps reclassified into two categories, the focus category (barren land) and the background. (C) Simulated evolution of the site. (D) Complementary CDFs of patch size for the patterns in row B (blue) and row C (orange). (E) Complementary CDFs of patch area for the patterns in row B (blue) and row C (orange). The numbers above observed and simulated patterns are the number of barren land cells and the pattern’s texture parameter ⟨*s*_*i*_ *s* _*j*_⟩. Numbers below the simulated patterns are accuracy, recall, and precision.

The fourth row in Table 1 presents the best-fit values of *B* and *J* for each transition in the series depicted in Fig. 6. Notably, the values of *B* exhibit significant variability from one observation year to another, indicating an uneven external influence. One plausible explanation for this variability is the fluctuating demand for coal. Conversely, the values of *J* remain relatively constant during the observed period. Similarly to the previous examples, we observe a high visual similarity between the simulated and observed land series. Furthermore, a quantitative analysis reveals that the simulated and observed patterns are characterized by nearly identical values of *m* and ⟨*s*_*i*_ *s* _*j*_⟩.

The values of recall and precision exhibit variability, ranging from 36% to 51% for the first two transitions, approximately 60% for the subsequent four transitions, and 82% in the final transition. These values display an inverse correlation with the rate at which the pit shifts westward, with lower accuracy observed during faster shifts and higher accuracy during slower shifts. This outcome can be attributed to the limitations of the IM model, which lacks a preferred direction in its dynamics and therefore cannot shift the pattern of focus cells from their initial position. The discrepancies observed in the shapes of patch size and area distributions are primarily attributed to the model’s inability to form linear features, as discussed in the previous subsections.

## 4. Discussion

Our hypothesis in the introduction suggested that the kinetic Ising model can serve as a framework for understanding various land change processes. By employing a bare-bones change model, we can abstract the underlying dynamics of change from the specific real-world processes associated with thematic contexts and locations of different sites. Furthermore, this model allows us to quantify change not only in terms of the extent of the altered area but also in terms of the intensity of the process. We now examine whether our results support the hypothesis that the IM can effectively serve as an abstraction of land change processes. The summary of our evaluation is as follows:

1. **Visual similarity:** We observed a high degree of visual similarity between the simulated and observed land series in all four case studies, indicating that the IM captures the overall patterns and dynamics of land change accurately.
2. **Quantitative measures:** The simulated and observed patterns exhibited nearly identical values of key quantitative pattern measures, such as the composition imbalance index *m* and the texture index ⟨*s*_*i*_ *s* _*j*_⟩. Patch size distribution in simulated and observed patterns also match except for the largest patches. This shows that the IM can effectively reproduce temporal changes in the composition and texture of landscape pattern.
3. **Parameter correlations:** We found strong correlations between inferred values of parameter *B* and rate of change. This indicates that the IM successfully captures the temporal variability of landscape composition in response to changing external forces.
4. **Model limitations:** They are some discrepancies between modeled and observed land change. The IM could not reproduce linear features, pattern shifts, and formation of large new patches.

Our findings provide compelling evidence that the fundamental principles of the IM, external influences and internal coupling between neighboring cells, are crucial factors driving various observed land change phenomena. This insight constitutes the primary original contribution of our research. By successfully demonstrating the IM’s capability to accurately capture the dynamics of land change across different thematic contexts and locations, we highlight that the specific intricacies of these underlying mechanisms may not be the determining factors in shaping the magnitude and nature of change. Instead, it is the interplay between external forcing and short-range interactions, regardless of their origin, that drives the observed patterns of change in land.

As mentioned in the introduction, the aim of this work was not to apply the IM model for land change prediction. However, our analysis of the four examples has demonstrated that the IM has the capability to predict the spatial arrangement of future patterns given the initial conditions, the future bulk composition, and an assumed constant value of *J*. The requirement of knowing the future bulk composition is common to all spatially-explicit models, including empirical, mechanistic, and agent-based models, as they rely on understanding the extent of change (referred to as a scenario, see, for example, Zhou et al. (2020)) in order to make predictions about future spatial patterns. As a cell-level predictor, the IM has shown reasonable accuracy for predicting reforestation and crop expansion sites, but lower accuracy for wetlands and mining sites. It is important to note that the accuracy of empirical predictors on wetlands and mining sites is also unclear, as these sites pose unique challenges for modeling.

Finally, it is important to highlight that our use of the IM model deviates from its conventional application. Typically, the IM is employed to explore the relationship between the equilibrium pattern characteristics and temperature, as well as to investigate critical phenomena. In our study, however, we deviate from these traditional uses. Our focus lies not on equilibrium patterns, temperature dependencies, or phase transitions. Instead, we employ a kinetic IM to simulate the evolution of a landscape, transitioning from a specific nonequilibrium pattern to another specific non-equilibrium pattern. Our main objective is to determine the optimal values of *J* and *B* that facilitate this transition within a given time frame.

In the examples we investigated, the landscapes were observed to be far from the critical point and away from equilibrium. Notably, the patch size distributions observed in these landscapes do not follow the expected power-law behavior of nearly critical patterns, except possibly for the wetlands patterns from 2013 to 2019 (Palmieri and Jensen, 2020; Pascual and Guichard, 2005). If we were to observe a landscape undergoing a “phase transition,” such as a transition from the background category to the focus category, our IM model would be capable of simulating such dynamics.

## References

Acheson, J., 2008. Maine: On the cusp of the forest transition 67 (2), 125–136.

Bornholdt, S., Wagner, F., 2002. Stability of money: phase transitions in an Ising economy. Physica A: Statistical Mechanics and its Applications 316 (1-4), 453–468.

Brandt, M. J., Sleegers, W. W., 2021. Evaluating belief system networks as a theory of political belief system dynamics. Personality and Social Psychology Review 25 (2), 159–185.

Brush, S. G., 1967. History of the Lenz-Ising model. Reviews of modern physics 39 (4), 883.

Camacho Olmedo, M., Mas, J., Paegelow, M., 2018. The simulation stage in LUCC modeling. In: Geomatic approaches for modeling land change scenarios. Springer, pp. 27–51.

Chen, G., Li, X., Liu, X., Chen, Y., Liang, X., Leng, J., Xu, X., Liao, W., Qiu, Y., Wu, Q., Huang, K., 2020. Global projections of future urban land expansion under shared socioeconomic pathways 11.

Cihlar, J., 2000. Land cover mapping of large areas from satellites: status and research priorities 21 (6-7), 1093–1114.

Cipra, B. A., 1987. An introduction to the Ising model. The American Mathematical Monthly 94 (10), 937–959.

Cramer, A. O., Van Borkulo, C. D., Giltay, E. J., Van Der Maas, H. L., Kendler, K. S., Scheffer, M., Borsboom, D., 2016. Major depression as a complex dynamic system. PloS one 11 (12), e0167490.

D’Autilia, R., Andrianaivo, L. N., Troiani, A., 2021. Parallel Simulation of Two-Dimensional Ising Models Using Probabilistic Cellular Automata 184 (1), 1–22.

Dewitz, J., 2021. National Land Cover Database (NLCD) 2019 Products (ver. 2.0): U.S. Geological Survey data release. URL https://doi.org/10.5066/P9KZCM54

Fajardo, A., Velázquez, E., 2021. Fine-scale spatial associations between functional traits and tree growth 130 (11), 1988–2000.

Gardner, R. H., Milne, B. T., Turnei, M. G., O’Neill, R. V., 1987. Neutral models for the analysis of broad-scale landscape pattern 1 (1), 19–28.

Gaucherel, C., Houet, T., 2009. Preface to the selected papers on spatially explicit landscape modelling: current practices and challenges 220, 3477–3480.

Gaucherel, C., Houllier, F., Auclair, D., Houet, T., 2014. Dynamic landscape modelling: the quest for a unifying theory 8(2), 5–31.

Glauber, R. J., 1963. Time-Dependent Statistics of the Ising Model 4, 294–307.

Gustafson, E. J., 2013. When relationships estimated in the past cannot be used to predict the future: using mechanistic models to predict landscape ecological dynamics in a changing world 28, 1429– 1437.

Hansen, M. C., Potapov, P. V., Moore, R., Hancher, M., Turubanova, S., Tyukavina, A., Thau, D., Stehman, S. V., Goetz, S. J., Loveland, T. R., Kommareddy, A., Egorov, A., Chini, L., Justice, C. O., Townshend, J. R., 2013. High-resolution global maps of 21st-century forest cover change. Science 342, 850–853.

Husmann, K., Lange, A., Spiegel, E., 2017. The R Package optimization: Flexible Global Optimization with Simulated-Annealing.

Ising, E., 1924. Beitrag zur theorie des ferro-und paramagnetismus. Ph.D. thesis, Grefe & Tiedemann.

Jamal, S., Ahmad, W. S., 2020. Assessing land use land cover dynamics of wetland ecosystems using Landsat satellite data 2 (11), 1–24.

Jamaludin, J., Alban, J. D. T. D., Carrasco, L. R., Webb, E. L., 2022. Spatiotemporal analysis of deforestation patterns and drivers reveals emergent threats to tropical forest landscapes 17.

Katori, M., Kizaki, S., Terui, Y., Kubo, T., 1998. Forest dynamics with canopy gap expansion and stochastic Ising model 6 (01), 81–86.

Kirkpatrick, S., Gelatt Jr, C. D., Vecchi, M. P., 1983. Optimization by simulated annealing 220 (4598), 671–680.

Kizaki, S., Katori, M., 1999. Analysis of canopy-gap structures of forests by Ising-Gibbs states-equilibrium and scaling property of real forests. Journal of the Physical Society of Japan 68 (8), 2553– 2560.

Koenig, W. D., 1999. Spatial autocorrelation of ecological phenomena 14 (1), 22–26.

Kumar, V., Agrawal, S., 2022. Urban modelling and forecasting of landuse using SLEUTH model, 1–20.

Lark, T. J., Spawn, S. A., Bougie, M., Gibbs, H. K., 2020. Cropland expansion in the United States produces marginal yields at high costs to wildlife 11 (1), 4295.

Li, W., MacBean, N., Ciais, P., Defourny, P., Lamarche, C., Bontemps, S., Houghton, R. A., Peng, S., 2018. Gross and net land cover changes in the main plant functional types derived from the annual ESA CCI land cover maps (1992–2015) 10, 219–234.

Liu, T., Yang, X., 2015. Land Change Modeling: Status and Challenges. In: Monitoring and Modeling of Global Changes: A Geomatics Perspective. Springer, Dordrecht., pp. 3–16.

Ma, Y.-P., Sudakov, I., Strong, C., Golden, K. M., 2019. Ising model for melt ponds on Arctic sea ice. New Journal of Physics 21 (6), 063029.

Majewski, J., Li, H., Ott, J., 2001. The Ising model in physics and statistical genetics. The American Journal of Human Genetics 69 (4), 853–862.

Metropolis, N., Rosenbluth, A. W., Rosenbluth, M. N., Teller, A. H., Teller, E., 1953. Equation of state calculations by fast computing machines. The journal of chemical physics 21 (6), 1087–1092.

Nareddy, V. R., Machta, J., Abbott, K. C., Esmaeili, S., Hastings, A., 2020. “dynamical ising model of spatially coupled ecological oscillators” 17 (171), 20200571.

Noble, A. E., Rosenstock, T. S., Brown, P. H., Machta, J., Hastings, A., 2018. Spatial patterns of tree yield explained by endogenous forces through a correspondence between the Ising model and ecology. Proceedings of the National Academy of Sciences 115 (8), 1825–1830.

Nowosad, J., 2023. spatialising: Ising Model for Spatial Data. R package version 0.4.0. URL https://github.com/Nowosad/spatialising

Nowosad, J., Stepinski, T. F., Netzel, P., 2019. Global assessment and mapping of changes in mesoscale landscapes: 1992-2015 78, 332–340.

Onsager, L., 1944. Crystal statistics. I. A two-dimensional model with an order-disorder transition. Physical Review 65 (3-4), 117.

Palmieri, L., Jensen, H. J., 2020. Investigating critical systems via the distribution of correlation lengths 2 (1), 013199.

Pascual, M., Guichard, F., 2005. Criticality and disturbance in spatial ecological systems 20 (2), 88–95.

Procacci, A., Scoppola, B., Scoppola, E., 2016. Probabilistic cellular automata for low-temperature 2-d Ising model 165 (6), 991–1005.

Rastetter, E. B., Aber, J. D., Peters, D. P., Ojima, D. S., Burke, I. C., 2003. Using mechanistic models to scale ecological processes across space and time 53 (1), 68–76.

Rosenfeld, A., Pfaltz, J. L., 1966. Sequential operations in digital picture processing 13 (4), 471–494.

Scanlon, T. M., Caylor, K. K., Levin, S. A., Rodriguez-Iturbe, I., 2007. Positive feedbacks promote power-law clustering of Kalahari vegetation. Nature 449 (7159), 209–212.

Scheller, R. M., Domingo, J. B., Sturtevant, B. R., Williams, J., Rudy, A., Gustafson, E. J., Mladenoff, D. J., 2007. Design, development, and application of LANDIS-II, a spatial landscape simulation model with flexible temporal and spatial resolution 201, 409– 419.

Schlicht, R., Iwasa, Y., 2006. Deviation from power law, spatial data of forest canopy gaps, and three lattice models. Ecological modelling 198 (3-4), 399–408.

Song, X.-P., Hansen, M. C., Stehman, S. V., Potapov, P. V., Tyukavina, A., Vermote, E. F., Townshend, J. R., 2018. Global land change from 1982 to 2016 560 (7720), 639–643.

Stauffer, D., 2008. Social applications of two-dimensional Ising models. American Journal of Physics 76 (4), 470–473.

Stauffer, D., Solomon, S., 2007. Applications of physics and mathematics to social science.

Stepinski, T. F., 2023. Spatially explicit simulation of deforestation using the ising-like neutral model. Environmental Research: Ecology.

Tomasella, J., Vieira, R. M. S. P., Barbosa, A. A., Rodriguez, D. A., de Oliveira Santana, M., Sestini, M. F., 2018. Desertification trends in the Northeast of Brazil over the period 2000–2016 73, 197–206.

Turner, M. G., 1989. Landscape Ecology: The Effect of Pattern on Process 20, 171–197.

Valbuena, D., Verburg, P. H., Bregt, A. K., Ligtenberg, A., 2010. An agent-based approach to model land-use change at a regional scale 25, 185–199.

Veldkamp, A., Lambin, E. F., 2001. Predicting land-use change 85, 1–6.

Venter, O., Sanderson, E. W., Magrach, A., Allan, J. R., Beher, J., Jones, K. R., Possingham, H. P., Laurance, W. F., Wood, P., Fekete, B. M., et al., 2016. Sixteen years of change in the global terrestrial human footprint and implications for biodiversity conservation 7 (1), 1–11.

Winkler, K., Fuchs, R., Rounsevell, M., Herold, M., 2021. Global land use changes are four times greater than previously estimated 12, 2501.

Zhou, L., Dang, X., Sun, Q., Wang, S., 2020. Multi-scenario simulation of urban land change in Shanghai by random forest and CA-Markov model 55, 102045.

